# Soil microenvironment and microbial community composition jointly regulate carbon accrual in agricultural soils

**DOI:** 10.64898/2026.09.18.752463

**Authors:** Lisa Cole, Tao Li, Tim Goodall, Rob Griffiths, Wolfgang Wanek, Emily MacDonald, Nicole Cochrane, Eric Paterson, Cécile Gubry-Rangin, Ashish A. Malik

## Abstract

Much of the persistent soil organic carbon (SOC) pool is microbial in origin: microorganisms process plant carbon into biomass and their necromass can associate with mineral surfaces, contributing to long-term carbon persistence. This has generated interest in microbial interventions such as inoculation to restore carbon in degraded croplands, yet their efficacy remains uncertain. The uncertainty reflects a more fundamental, unresolved question: are microbially-mediated carbon transformations governed principally by the composition of the microbial community or by the soil microenvironment? To disentangle these drivers, we conducted a reciprocal microbial community transplant experiment, introducing communities of contrasting origin from locally adjacent grassland and cropland soils into sterilised grassland and cropland soils, and incubating them for eight months with regular organic inputs. This design decouples the inoculum from the microenvironment, allowing their individual and interactive contributions to be quantified. Fungal assembly was influenced more by the inoculum, consistent with dispersal limitation, whereas bacterial assembly was governed more by the microenvironment, consistent with environmental selection. Despite receiving the same organic carbon inputs, grassland and cropland recipient soils showed distinct SOC trajectories, indicating that the soil microenvironment strongly constrained net carbon retention. Within this constraint, community composition also mattered: introducing grassland rather than cropland communities increased fungal diversity and fungal necromass and led to better SOC outcomes, expressed as net gain or reduced loss. These results show that SOC accrual emerges from interactions between the soil microenvironment and microbial community composition, with fungal community assembly particularly associated with necromass accumulation and carbon retention. They highlight the need to consider both soil conditions and microbial community composition when developing strategies to enhance SOC accrual.

**Significance Statement:** Restoring carbon in degraded agricultural soils offers an opportunity to mitigate climate change while improving soil health and supporting food security. Because much of the persistent soil carbon pool is microbial in origin, there is growing interest in using microbial interventions to enhance carbon retention in degraded farmland. Yet it remains unclear whether soil carbon outcomes are governed more by which microbes are present or by the soil conditions in which they operate. Using a reciprocal transplant experiment that exchanged microbial communities between grassland and cropland soils, we show that soil conditions strongly constrain carbon retention, while community composition—particularly the diversity of fungi—modulates the outcome within that constraint. Preserving and rebuilding soil carbon will therefore require consideration of both soil conditions and its microbial community.

## Introduction

Soils are the largest terrestrial carbon reservoir. Agricultural intensification has led to substantial losses of soil carbon; some estimates suggest that 133 Gt has been lost historically (Sanderman et al., 2017). Rebuilding SOC in agricultural soils, therefore, offers the opportunity to mitigate climate change while restoring soil health. For a long time, the persistence of SOC was attributed to the intrinsic molecular recalcitrance of plant compounds. This view has been overturned as persistence is now understood as an emergent system-level property, governed by organo-mineral associations and physical protection rather than by molecular structure alone (Schmidt et al., 2011; Lehmann and Kleber, 2015). Central to this reconceptualization is the recognition that much of the persistent SOC pool is microbial in origin (Liang et al., 2017; Malik, 2026), although plant-derived compounds may also be stabilised directly (Angst et al., 2021; Elias et al., 2024).

Soil microorganisms decompose plant organic matter and release a large fraction of the acquired carbon through respiration, while the remainder contributes to living biomass and, following turnover, to dead cellular residues termed necromass (Kallenbach et al., 2016; Sokol et al., 2022). These microbial residues can associate efficiently with soil minerals, and contribute substantially to the formation of relatively persistent mineral-associated organic matter (Kleber et al., 2007, 2015; Lehmann and Kleber, 2015). How much carbon persists therefore depends both on the rate at which carbon is supplied and on how microorganisms partition acquired carbon between growth and respiration. This partitioning is captured by an aggregate, community-level trait – microbial carbon use efficiency or CUE, (Manzoni et al., 2018) – which generally correlates with soil organic carbon across environmental gradients (Malik et al., 2018, 2020; Feng et al., 2021; Mganga et al., 2022; Tao et al., 2023). Higher CUE favours greater biomass yield per unit carbon acquired, while substrate supply, microbial growth and turnover regulate rates of biomass production and necromass formation (Kästner et al., 2021; Whalen et al., 2024). Necromass accumulation provides an integrative record of its production, decomposition and stabilisation (Buckeridge et al., 2022). This places microbial community ecophysiology at the heart of soil carbon persistence. Managing the factors that govern microbial growth and necromass accumulation may therefore offer a route to greater soil carbon accrual.

The effect of microbial ecophysiology on soil carbon persistence could depend on two controlling factors that are not mutually exclusive. The first is the microbial community composition: the identity, diversity, and functional repertoire of the assembled taxa and the phenotypic expression of those functions together determine carbon transformations (Domeignoz-Horta et al., 2021). The second factor is the microenvironment: the mineralogy, aggregate structure, moisture, and resource availability that together determine both microbial activity and the capacity to retain microbial products (Wasner et al., 2024; Liu et al., 2026). In resource-rich, optimum-moisture soil environments such as those under low-intensity grasslands, microorganisms tend to have higher CUE, whereas in the resource-poor, drier and more poorly structured soils typical of intensively managed croplands, they invest proportionally more carbon in stress tolerance and resource acquisition, lowering CUE (Malik et al., 2018; Cole et al., 2024). The microenvironment may thus exert control both by acting as an environmental filter that selects for taxa possessing particular carbon-transformation traits and by modulating the phenotypic expression of the community that establishes.

Community composition is itself shaped by ecological assembly processes. Along with environmental selection, dispersal can also affect the composition of the established community (Vellend, 2010; Nemergut et al., 2013). If environmental filtering is the dominant control, communities introduced into a given soil should converge toward the resident state regardless of origin; if community identity dominates, they should instead retain a legacy of their source. These processes need not act equally across the microbiome: fungi and bacteria differ in dispersal ability, physiology, life history, cellular structure, and molecular composition, and may therefore be governed by different assembly processes and contribute differently to necromass formation (Sokol et al., 2022; Camenzind et al., 2023). Distinguishing these controls on soil carbon persistence in intact soils is difficult because community and microenvironment are inextricably linked. Isolating their individual and interactive contributions therefore requires experimentally decoupling the community from the environment in which it assembled.

The microbial contribution to persistent SOC has motivated proposals for microbial interventions, such as inoculation of degraded croplands with communities or consortia selected to enhance carbon accrual. Yet the efficacy of such interventions remains uncertain as introduced taxa often fail to establish, are outcompeted by resident taxa, or exert only transient effects on ecosystem function (Kaminsky et al., 2019; Mawarda et al., 2022). While there are technical obstacles in implementing such interventions, there is also a more fundamental knowledge gap concerning whether soil carbon outcomes are governed principally by community composition or by the soil microenvironment. Until this is resolved, such interventions cannot be designed on a rational basis.

Here we investigated whether microbial transformation of plant carbon into biomass and necromass, and its consequences for SOC accrual, are controlled more by which microorganisms are present or by the environment in which they operate. We hypothesise that carbon accrual is an emergent property of the soil system, shaped jointly by the composition of the microbial community and the soil microenvironment. To quantify their relative and interactive contributions, we conducted a reciprocal microbial community transplant experiment, introducing microbial communities from locally adjacent grassland and cropland soils into sterilised cropland and grassland soils. Sterilising the soils and reintroducing communities of contrasting origin allowed us to decouple the community from its environment. We incubated the soil systems for eight months while maintaining moisture and supplementing with plant-litter-derived dissolved organic carbon. At the end of the experiment, we characterised the assembled microbial communities and quantified associated carbon outcomes using microbial necromass and total SOC. Specifically, we tested whether inoculating cropland soils with a grassland community would enhance necromass accumulation and SOC retention, or whether the cropland soil microenvironment would constrain these outcomes regardless of the community introduced.

## Results

### Microbial community assembly and diversity

We first characterised the microbial communities that assembled in the experimental mesocosms eight months after inoculation of the sterilised soils, using amplicon sequencing of the 16S rRNA gene for prokaryotes (bacteria and archaea) and the ITS region for fungi. Assembly of both prokaryotic and fungal communities was shaped by soil properties and inoculum source (Fig. 1), but their relative importance differed between the two groups. Soil properties explained more of the variance in prokaryotic community composition (22% at Site 1, 32% at Site 2) than inoculum source did (19% and 16%, respectively). For fungi, the opposite pattern emerged: inoculum source explained more variance (29% at Site 1, 21% at Site 2) than soil properties (12% and 10%, respectively).

**Fig. 1.**
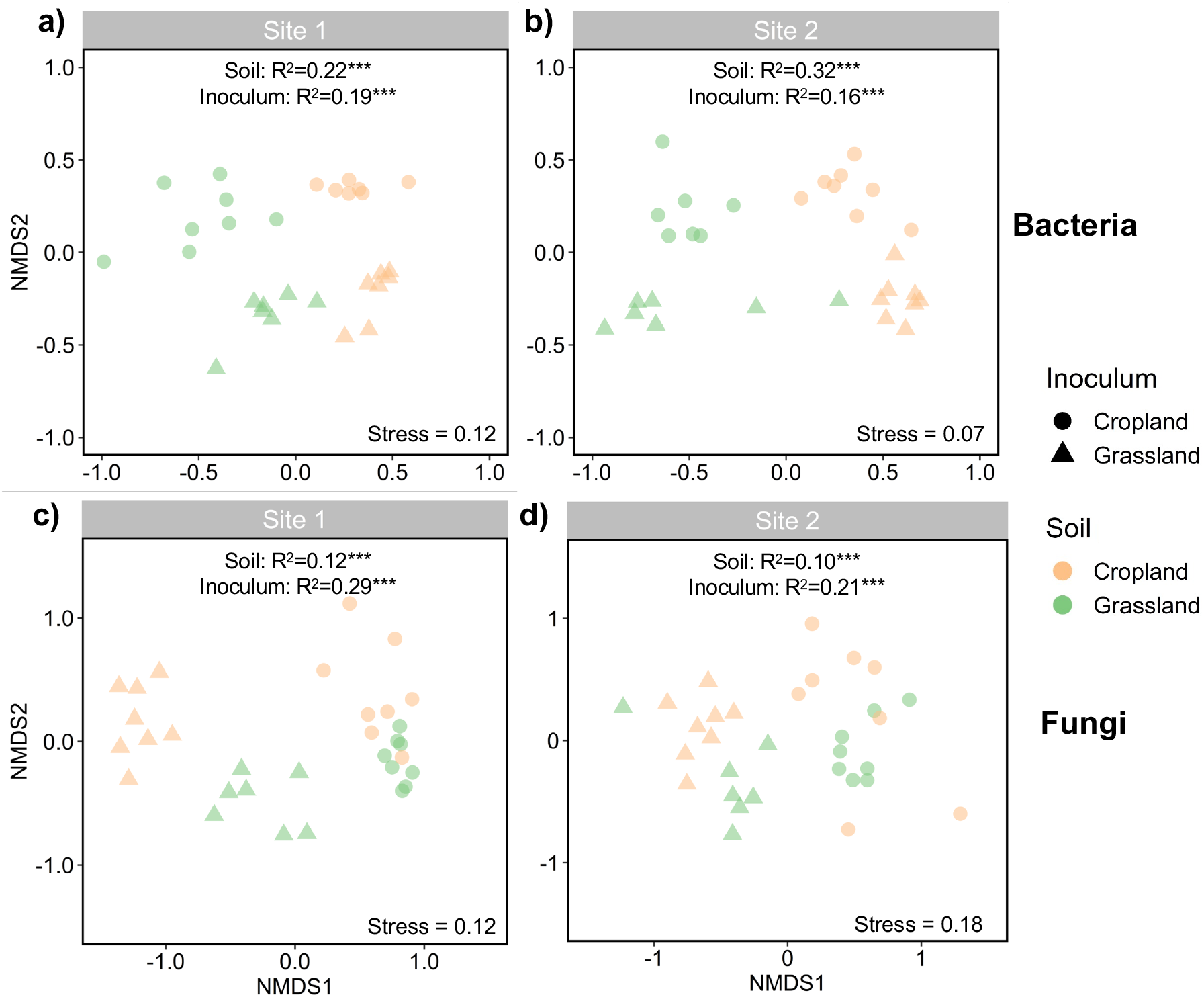
Composition of the assembled community. NMDS ordination for prokaryotic ASVs derived from 16S rRNA gene sequencing for Site 1 (a) and Site 2 (b), and for fungal ASVs derived from ITS sequencing for Site 1 (c) and Site 2 (d). PERMANOVA results by experimental factors, and NMDS stress scores are displayed in each plot; *P*<0.001***.

Prokaryotic alpha diversity (Shannon index) was largely similar among individual treatments in Tukey’s post hoc tests (Fig. 2a), although soil properties had a significant effect at Site 1 and inoculum source at Site 2 (ANOVA, *P* < 0.01 in both cases). Across treatments, prokaryotic alpha diversity tended to be higher when a given soil received the cropland rather than the grassland inoculum, and prokaryotic richness and evenness followed the same trend (Fig. S1). Fungal alpha diversity showed the opposite pattern, being significantly higher when a given soil received the grassland rather than the cropland inoculum (Fig. 2b, S1).

**Fig. 2.**
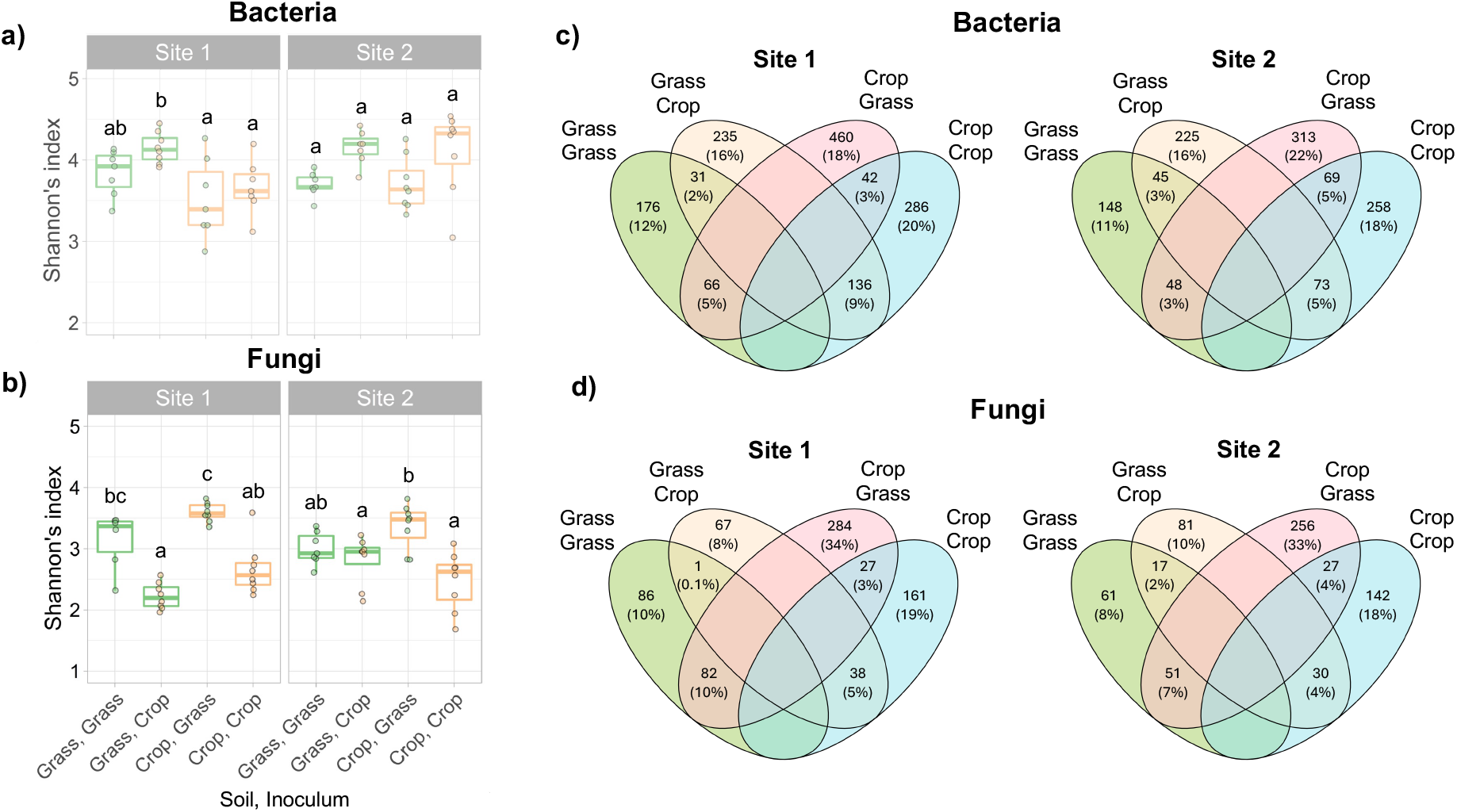
Diversity and unique taxa in the assembled community. Boxplot of Shannon’s diversity index for prokaryotic ASVs (a), and fungi (b). Letters show pairwise Tukey HSD test results at p < 0.05 analysed for each site separately; shared letters signify lack of statistical significance. Thick horizontal lines in the boxplots show median, boxes show interquartile ranges and vertical whiskers extend 1.5 times the inter quartile range. Venn diagrams represent unique and shared prokaryotic (c) and fungal (d) ASVs across treatments.

We next quantified the taxa unique to each treatment. For prokaryotes, the proportion of unique ASVs was similar across inoculum treatments within a given soil: grassland soils contained 11–12% unique ASVs with the grassland inoculum and 16% with the cropland inoculum, while cropland soils contained 18–20% and 18–22% unique ASVs with the cropland and grassland inocula, respectively (Fig. 2c). For fungi, inoculum source had a much stronger effect. Grassland soils contained 8–10% unique fungal ASVs under both inocula, whereas cropland soils with the cropland inoculum contained 18–19% unique fungal ASVs. In contrast, cropland soils receiving the grassland inoculum contained a substantially higher proportion of unique fungal ASVs (33–34%) and the highest fungal diversity of any treatment (Fig. 2d).

We then identified indicator taxa associated with the different soil–inoculum combinations (Fig. S2). For bacteria, more taxa were shared among mesocosms constructed from the same sterilised soil than among mesocosms receiving the same inoculum. Grassland soils selected for diverse taxa within the Actinobacteriota and Planctomycetota regardless of inoculum, although the grassland inoculum reinforced this pattern; a similar response occurred for Firmicutes, but only at Site 1. Cropland soils, by contrast, showed lower evenness and were dominated by a few highly abundant taxa within the Actinobacteriota (Micrococcales) and Proteobacteria (Burkholderiales), again largely regardless of inoculum. The only bacteria consistently selected with the cropland inoculum across soils belonged to the Bacteroidota, with an additional Firmicutes response at Site 2.

Fungi showed the opposite structuring: more taxa were shared among mesocosms receiving the same inoculum than among those constructed from the same soil, with few discernible taxa consistently associated with a given recipient soil. The grassland inoculum was associated with the greatest number of unique fungal taxa across both soils, mostly Ascomycota, with additional Mucoromycota at Site 1 and Mortierellomycota. A smaller set of abundant fungi was common to mesocosms receiving the cropland inoculum, comprising Ascomycota (Hypocreales) and, at Site 1, Mortierellomycota (Mortierellales).

The assembled communities remained distinct from those originally present in the non-sterilised field soils (Fig. S3-5), indicating that inoculation into sterilised soil did not fully re-establish the original communities within eight months.

### Soil respiration and microbial biomass

We monitored soil respiration regularly across the eight-month mesocosm incubation to track microbial activity. Following re-inoculation and rewetting, respiration showed a pronounced initial pulse and declined toward a relatively stable baseline after approximately 45 days (Fig. S6), before regular additions of plant-litter-derived DOC began on day 50. Cumulative respiration across the incubation was greater from grassland than cropland recipient soils at both sites (ANOVA, soil properties P < 0.001 at both sites). Respiration was lower when soils received the cropland inoculum rather than the grassland inoculum, although this inoculum effect was significant only at Site 1 (ANOVA, inoculum P < 0.001).

Microbial biomass, estimated from total DNA yield at the end of the experiment, was significantly higher in grassland than cropland recipient soils at both sites (Fig. S7; ANOVA, soil properties P < 0.001 at both sites), whereas no significant inoculum effect was detected. DNA-derived microbial biomass nevertheless showed substantial variation among replicate mesocosms.

### Microbial necromass at the end of incubation

We estimated microbial necromass at the end of the incubation using amino sugar biomarkers, deriving bacterial- and fungal-derived necromass from muramic acid (bacterial) and glucosamine (bacterial and fungal). Both bacterial and fungal necromass were greater overall in grassland than cropland recipient soils at both sites (Fig. 3; mixed effects model, soil properties P < 0.001 at both sites).

**Fig. 3.**
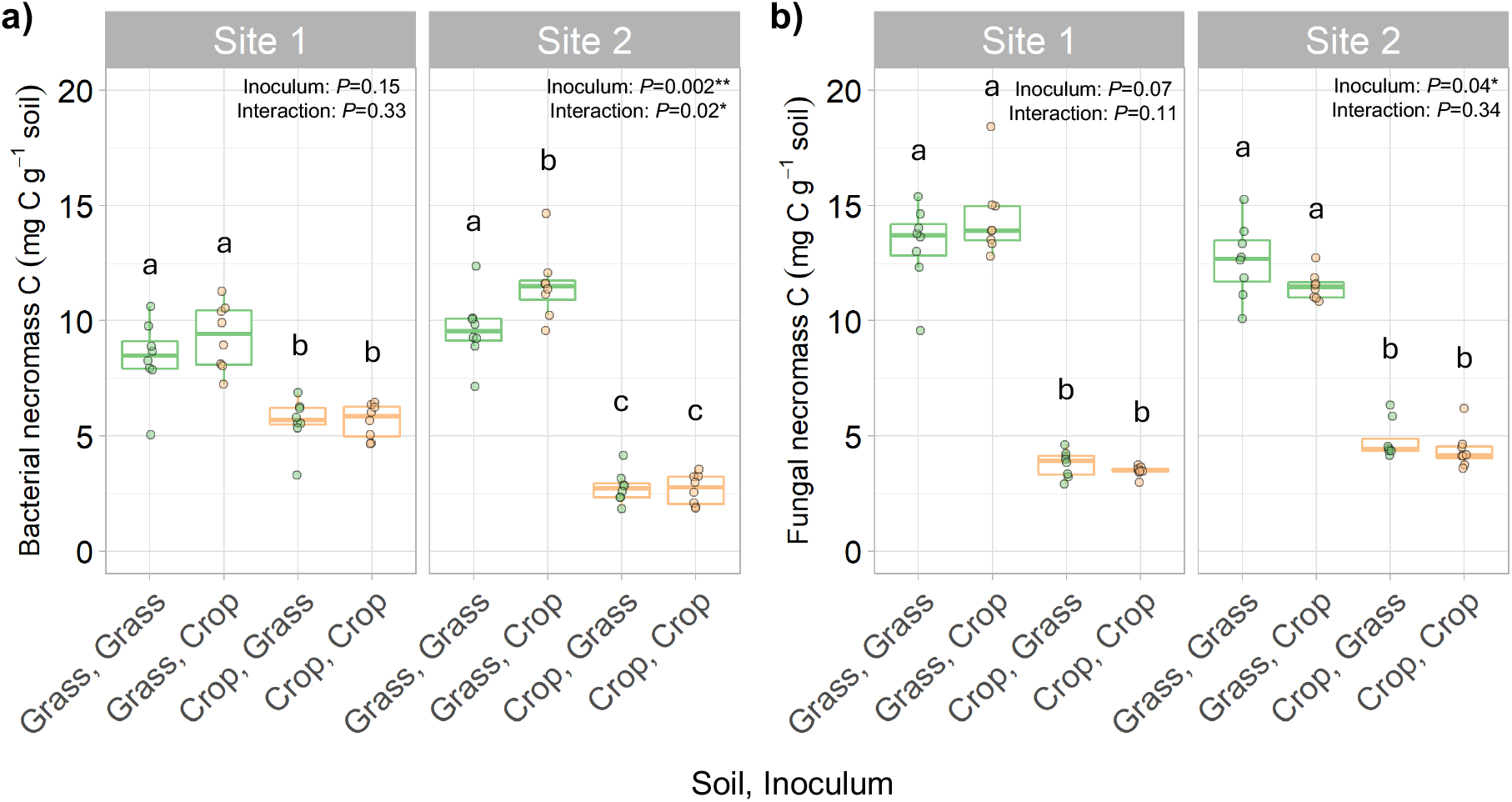
Microbial necromass at the end of the experiment. Bacterial (a) and fungal (b) necromass at the end of the experiment. Letters show pairwise Tukey HSD test results at p < 0.05 analysed for each site separately; shared letters signify lack of statistical significance. Thick horizontal lines in the boxplots show median, boxes show interquartile ranges and vertical whiskers extend 1.5 times the inter quartile range. *P* values show results of mixed effects models assessing the effect of soil properties, inoculum source and their interactions \**P* < 0.1, \*\**P* < 0.01. Soil properties were always significant (*P* < 0.001) and are not displayed.

For bacterial necromass, the inoculum-source effect was significant only at Site 2, where grassland soils receiving the cropland inoculum contained more bacterial necromass than those receiving the grassland inoculum. The soil properties × inoculum interaction was also significant at Site 2, indicating that the inoculum effect depended on soil properties (mixed effects model; Site 1: inoculum P = 0.15, interaction P = 0.33; Site 2: inoculum P = 0.002, interaction P = 0.02).

For fungal necromass, the inoculum-source effect was significant at Site 2 and marginal at Site 1, with fungal necromass generally higher under the grassland than the cropland inoculum. The soil properties × inoculum interaction was not significant at either site (mixed effects model; Site 1: inoculum P = 0.07, interaction P = 0.11; Site 2: inoculum P = 0.04, interaction P = 0.34).

### Change in SOC over the experimental period

Final SOC was higher in grassland than cropland recipient soils, reflecting long-term differences in land management (Fig. 4a; mixed effects model, soil properties P < 0.001). Inoculum source also affected final SOC: at both sites, soils receiving the grassland rather than the cropland inoculum had higher mean SOC. At Site 1, final SOC in sterilised cropland soil was significantly higher when it received the grassland rather than the cropland inoculum

**Fig. 4.**
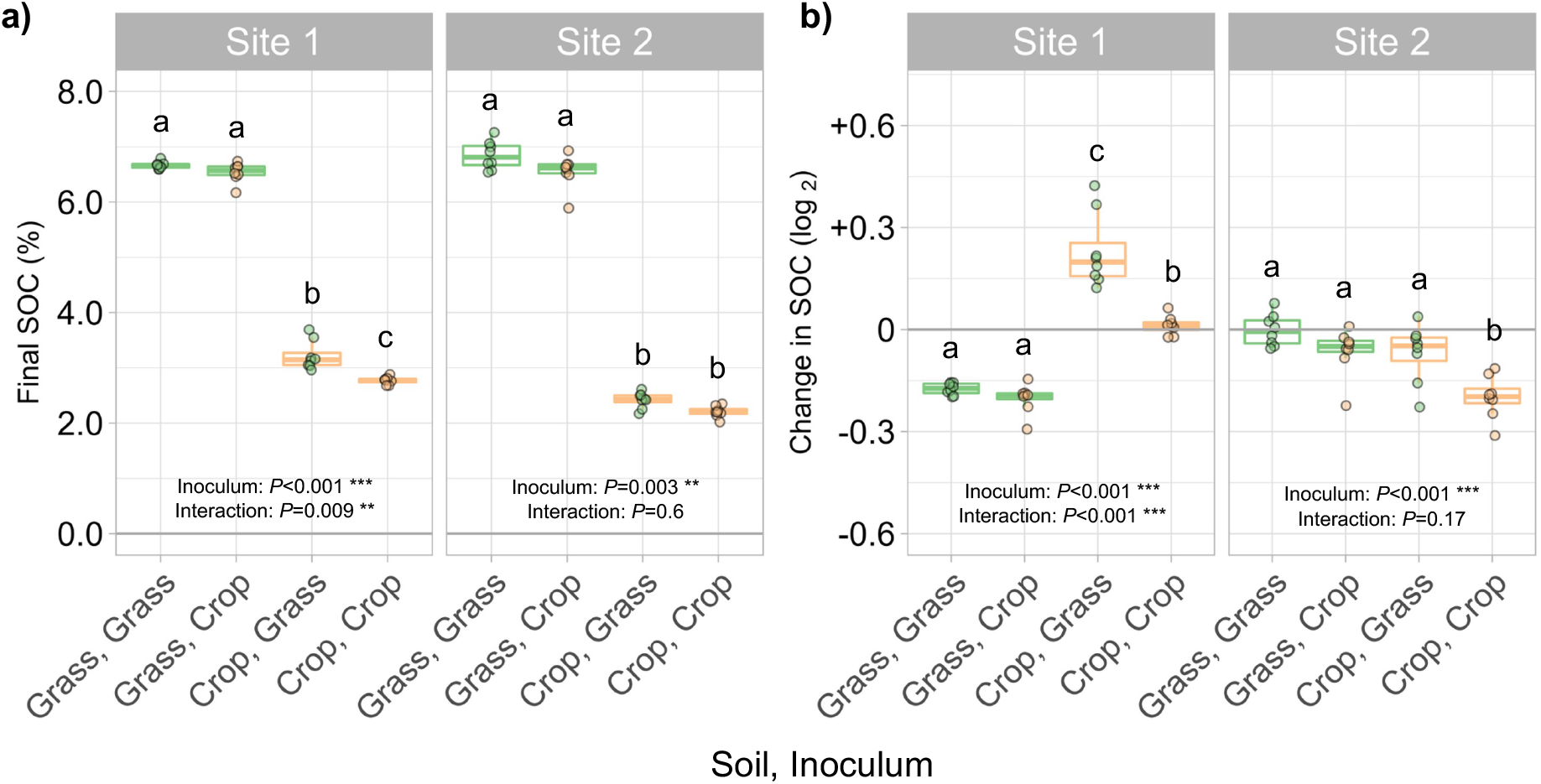
Soil carbon change at the end of the experiment. (a) Final SOC soils across the treatments and (b) the change in SOC over the experimental period expressed as the ratio of final and initial SOC content for each soil mesocosm. Letters show pairwise Tukey HSD test results at p < 0.05 analysed for each site separately; shared letters signify lack of statistical significance. Thick horizontal lines in the boxplots show median, boxes show interquartile ranges and vertical whiskers extend 1.5 times the inter quartile range. *P* values show results of mixed effects models assessing the effect of soil properties, inoculum source and their interactions. Soil properties were always significant (*P* < 0.001) and are not displayed.

SOC change over the eight-month incubation showed a similar treatment pattern (Fig. 4b; mixed effects model; Site 1: inoculum P < 0.001, interaction P < 0.001; Site 2: inoculum P < 0.001, interaction P = 0.17). At Site 2, cropland soil lost significantly more SOC when inoculated with its own community than when receiving the grassland inoculum (Fig. 4b). Sterilised grassland soils at both sites lost SOC, with a tendency toward greater loss under the cropland than the grassland inoculum. Overall, cropland soils receiving the grassland inoculum showed more favourable SOC trajectories than those receiving the cropland inoculum, expressed as either net SOC gain at Site 1 or reduced SOC loss at Site 2.

## Discussion

### Contrasting controls on prokaryotic and fungal community assembly

Community assembly following inoculation revealed contrasting controls for the major microbial groups. Prokaryotic communities were more strongly structured by recipient soil than by inoculum source, whereas fungal communities showed the opposite pattern, with a stronger legacy of inoculum origin. This suggests that local soil conditions exerted a stronger environmental filtering effect on prokaryotic communities, while fungal assembly remained more dependent on the composition of the introduced community. The clearest illustration was that grassland-derived inoculum added to sterilised cropland soil produced the highest fungal diversity of any treatment, exceeding even sterilised grassland soil that received its own grassland inoculum. This stronger inoculum dependence in fungi is consistent with broader evidence that fungal communities can be more constrained by dispersal and propagule supply than bacterial communities (Chen et al., 2020; Zhang et al., 2021; Gill et al., 2022), although dispersal itself was not directly manipulated here. Differences in dispersal mode, propagule size, hyphal growth, and sensitivity to disturbance (Cho et al., 2017; Osburn et al., 2019) may contribute to the stronger source-community legacy observed for fungi. On the contrary, diverse bacterial taxa appear to be widely present, if at low abundance, and rise to dominance where local conditions favour their growth.

### Grassland-derived fungi improve SOC outcomes in cropland soil

The contrasting responses of prokaryotic and fungal communities were accompanied by different SOC trajectories over the incubation. In cropland soils, addition of the grassland-derived inoculum resulted in more favourable SOC outcomes than addition of the cropland inoculum: SOC increased at Site 1 and SOC losses were reduced at Site 2. These treatments also supported greater fungal diversity and generally higher fungal necromass, suggesting that community composition contributed to differences in C retention within the constraints imposed by the recipient soil microenvironment. This indicates that community composition and the traits it encodes can promote necromass production and SOC accrual, consistent with previous evidence linking microbial community traits, including carbon-use efficiency, to SOC persistence (Malik et al., 2018; Tao et al., 2023). Fungal-dominated communities may contribute through greater biomass production per unit C acquired and through the formation of necromass that can become associated with mineral surfaces (Kallenbach et al., 2016; Camenzind et al., 2023), although these mechanisms were not directly quantified here. By contrast, SOC changed little when the grassland community was reintroduced into grassland soil at either site, suggesting that the effect of inoculum source depended strongly on the recipient soil environment. The results support our hypothesis that carbon accrual is an emergent property of both community composition and the soil physico-chemical environment.

### SOC losses were greatest under the cropland inoculum

Despite the SOC gain in cropland soil receiving grassland inoculum at Site 1, most mesocosms showed little net change or lost SOC over the incubation. These losses may reflect enhanced decomposition of pre-existing SOC following sterilisation and rewetting and in response to repeated additions of plant-litter-derived DOC. Because the origin of respired C was not traced isotopically, however, the relative contributions of added DOC and native SOC mineralisation cannot be resolved. SOC losses were generally greatest where the cropland inoculum was introduced, coinciding with lower fungal necromass, suggesting a potential link between fungal community composition, necromass accumulation, and net carbon retention. At Site 2, cropland soil lost less SOC under the grassland than the cropland inoculum, again coinciding with greater fungal richness and fungal necromass. Across both sites, therefore, cropland soils receiving the grassland community showed more favourable C outcomes—either net SOC gain or reduced SOC loss—than those receiving their own community. Although the processes responsible for these differences cannot yet be separated, the consistent inoculum effect demonstrates that microbial community composition can modify SOC trajectories in degraded cropland soils. This provides proof of principle that microbial interventions may influence soil C retention, while the specific mechanisms and their field applicability remain to be established.

### Fungal necromass tracked SOC outcomes

Fungal and bacterial necromass pools showed divergent relationships with SOC change. Treatments and sites with higher fungal necromass tended to show more favourable SOC trajectories, either gaining carbon or losing less carbon over the incubation. This was not the case for bacterial necromass, where the cropland inoculum increased bacterial necromass at Site 2 without a corresponding improvement in SOC change. This contrast suggests that fungal and bacterial necromass may contribute differently to net SOC retention in these soils. Several mechanisms could contribute. Fungal cell-wall compounds such as chitin and melanin may differ from bacterial residues in their susceptibility to decomposition and interactions with soil minerals, while hyphal growth may increase contact with mineral surfaces (Camenzind et al., 2023; Beidler et al., 2024; Cong et al., 2025; Etesami, 2026). Conversely, bacterial necromass may be recycled more rapidly within the microbial community (Hu et al., 2018; Buckeridge et al., 2022). These mechanisms remain speculative here, however, because neither mineral association nor necromass turnover was measured directly.

Amino sugars represent standing necromass pools rather than fluxes, so the present data demonstrate an association between fungal necromass, and SOC change rather than direct differences in production, stabilization, or persistence. The relative turnover and retention of bacterial and fungal necromass remain uncertain (e.g., Buckeridge et al., 2020), and resolving these processes will require isotopic or time-resolved approaches.

### Saprotrophic fungi as a route to carbon restoration

Grassland-derived inoculum established diverse, functionally distinct fungal assemblages in cropland soil, accompanied by greater fungal necromass. Together with the association between fungal necromass and SOC change described above, this positions fungi as central to microbially-mediated soil restoration, consistent with evidence that fungal consortia can accelerate SOC accrual and successional processes (Emilia Hannula and Morriën, 2022; de Goede et al., 2025).

Soil inoculation is already used to return reclaimed agricultural land to grassland (Srivastava and Singh, 2022; Badger Hanson and Docherty, 2023), typically with mycorrhizal fungi as target consortia to accelerate the establishment of native plant communities (Neuenkamp et al., 2019; Koziol et al., 2022). In such studies, SOC gains are usually a secondary benefit that emerges once vegetation establishes, through reduced disturbance, increased plant inputs and associated changes in microbial CUE (Domeignoz-Horta et al., 2024). Our plant-free system shows that this need not be the case: saprotrophic fungi—including Hypocreales, Pleosporales, Xylariales, and Mortierellales— were associated with greater fungal necromass and more favourable SOC outcomes even in the absence of living plants. This highlights that traits beyond CUE, such as hyphal growth morphology, substrate processing, and necromass chemistry, that may influence how rapidly labile C is converted into longer-lived microbial residues and retained in soil (Whalen et al., 2024). Whereas current bioinoculant development centres largely on plant-associated mutualists such as mycorrhizal fungi and plant-growth-promoting bacteria, our results identify saprotrophic fungi as an under-explored target for enhancing SOC retention in degraded croplands.

### Implications for microbial interventions in agroecosystems

Our results show that microbial inoculation can influence SOC trajectories, but that its effects depend strongly on the recipient soil environment. Grassland-derived inocula consistently produced more favourable SOC outcomes in cropland soils, while community assembly differed in the relative importance of inoculum source and local soil conditions for fungi and prokaryotes. Together, these findings suggest contrasting controls on microbial establishment and persistence, with potential consequences for how fungal- and bacterial-mediated C processes respond to management and future biological interventions.

Field outcomes are likely to be more variable than those observed in our sterilised mesocosms. In intact soils, resident communities may resist establishment of introduced taxa, while abiotic stress, plant–microbe interactions, and management history will further influence persistence and function (Liu et al., 2022; Mawarda et al., 2022). Our conclusions are also based on two paired sites, and the specific mechanisms underlying the contrasting SOC responses remain unresolved. Long-term field experiments will therefore be required to test whether microbial inoculation can reproducibly alter community assembly, necromass formation, and SOC retention under realistic agricultural conditions (Sessitsch et al., 2019; O’Callaghan et al., 2022; Robinson et al., 2023). Restoring carbon in degraded croplands will ultimately require integrating soil management with microbial interventions to advance carbon sequestration, sustainable agriculture, and ecosystem restoration.

## Material and methods

### Sites and soil sampling

To investigate how microbial community composition and traits influence soil carbon dynamics under contrasting land-use intensity, we established a mesocosm experiment using soils collected from two UK sites previously characterised as part of a landscape-scale study (Malik et al., 2018). Site 1, at Knapwell, Cambridgeshire, comprised a diverse, herb-rich, historically uncultivated grassland (52°14’40.9”N, 0°02’59.5”W) that was grazed by sheep at the time of sampling, adjacent to cropland (52°14’46.6”N, 0°03’01.4”W) planted with spring barley (*Hordeum vulgare* L.). Site 2, at Wittenham, Oxfordshire, comprised a historically undisturbed, diverse grassland (51°37′40″N, 001°10′58″W) grazed by cattle, adjacent to cropland (51°37′38″N, 001°10′58″W) planted with broad beans (*Vicia faba* L.). Eight soil cores (3 cm diameter, 10 cm depth) were collected from each land-use type on 11^th^ and 12^th^ May 2022.

Following collection, soils were stored at 4°C and sieved to 4 mm to remove coarse material while minimizing disruption of soil aggregates. Subsamples from five cores per location were used to characterise soil physicochemical properties, including soil moisture and pH in water. The remaining soil was combined to generate a composite sample representative of each site × land-use combination.

A portion of the composite soil was dried at 30°C for 48 h and subsequently γ-irradiated (34.6-38.9 kGy, Steri-Ast Bradford, UK) to remove the resident microbiome while minimizing disruption of soil physical structure (McNamara et al., 2003; Merino-Martín et al., 2021). The remaining fresh composite soil was maintained at 4°C and used as a source of microbial inoculum. SOC content of the γ-irradiated soils was measured as described below.

### Experimental design, mesocosm construction, inoculation and maintenance

A reciprocal microbial transplant experiment comprising 64 mesocosms was established in October 2022 and maintained for 229 days (∼8 months). At each of the two sites, the experiment followed a 2 × 2 factorial design in which γ-irradiated grassland or cropland recipient soils were inoculated with microbial communities derived from either grassland or cropland soil. This yielded four treatment combinations: (1) grassland soil with grassland inoculum, (2) grassland soil with cropland inoculum, (3) cropland soil with grassland inoculum, and (4) cropland soil with cropland inoculum. Each treatment was replicated eight times at each site, yielding 64 mesocosms in total (2 sites × 2 recipient soils × 2 inoculum sources × 8 replicates).

Microbial inocula were prepared by extracting fresh composite donor soils with sterile artificial rainwater at a 1:2 soil-to-solution ratio (w:v) for 1 h on an orbital shaker, followed by filtration through Whatman No. 1 filter paper (Johnson et al., 2011). Mesocosms were constructed by adding 9 g dry γ-irradiated recipient soil to pre-weighed sterile 100-mL glass serum vials. Vials were stoppered with sterile sponge covered with sterile foil to permit gas exchange while minimizing cross-contamination among mesocosms.

During the first week of establishment, mesocosms received three additions of microbial wash and sterile artificial rainwater to restore gravimetric moisture contents approximating those of the corresponding field soils (Table 1: Site 1 grassland, 37%; Site 1 cropland, 22%; Site 2 grassland, 30%; Site 2 cropland, 22%). Each mesocosm received a total of 2.25 mL microbial wash together with 0.3 g fresh donor soil from the corresponding site and land-use type to facilitate successful community establishment.

Mesocosms were weighed after establishment and maintained at their target gravimetric moisture content by regular addition of sterile artificial rainwater and/or dissolved organic C solution. Two replicates of each treatment combination were randomly allocated to each of four trays, and tray positions were rotated weekly within the incubator to minimize positional effects. Mesocosms were incubated in the dark at 15 °C from days 0–171 and at 17 °C from days 172–229. The temperature was increased on day 172 to enhance soil drying and thereby permit further DOC additions intended to support microbial biomass prior to the final harvest on day 229.

### Dissolved organic carbon inputs

Because plants were absent from the mesocosms, plant-litter-derived dissolved organic carbon (DOC) was supplied as an external microbial C resource after the initial respiration pulse following rewetting and inoculation had declined toward a stable baseline. DOC solutions were prepared on two occasions from rinsed shoot material of ribwort plantain (*Plantago lanceolata* L.; 10 g), bird’s-foot trefoil (*Lotus corniculatus* L.; 5 g), and timothy hay (*Phleum pratense* L.; 10 g), dried at 40 °C and coarsely homogenised. Ribwort plantain and bird’s-foot trefoil were collected locally in Aberdeenshire and had been observed in the vegetation at Site 1, whereas timothy hay was commercially sourced.

The plant material was added to 500 mL sterile deionised water, heated to 100 °C, allowed to cool, and then shaken end-over-end for 24 h. The extract was filtered overnight through Whatman No. 1 filter paper (11 µm) and stored frozen at −20 °C until use. Before addition to the mesocosms, the DOC solution was thawed and sterilised by syringe filtration through a 0.45-µm membrane. Total organic carbon concentration was measured using a LABTOC TOC water analysis system (PPM Ltd, Kent, UK).

DOC was added on days 50, 75, 108, 131, 152, 168, 180, and 208. Each of the first four additions supplied 0.4 mg C g−^1^ soil, and each of the subsequent four additions supplied 0.31 mg C g−^1^ soil, giving a cumulative DOC-C input of 2.84 mg C g−^1^ soil over the incubation.

### Microbial respiration

Soil microbial respiration was monitored throughout the incubation as CO_2_ production, with measurements on days 1, 3, 8, 10, 15, 22, 29, 43, 57, 80, 102, 129, 150, 178, 199, and 224. For each measurement, mesocosms were sealed and incubated in the dark for 18 h at the prevailing incubation temperature (15 °C until day 171 and 17 °C thereafter). A 5-mL headspace gas sample was then collected into a 3-mL Exetainer and analysed within one week for CO_2_ concentration using an Agilent 6890 gas chromatograph (Agilent Technologies, Santa Clara, CA, USA). Respiration rates were expressed as C released per unit soil dry mass and time. Headspace CO2concentrations in ppm were converted to mg CO_2_ /m−^3^ and then to μgC-CO_2_ g−^1^ h−^1^ considering the volume of headspace, incubation time, and the dry weight of incubated soils.

### Destructive sampling of mesocosms

At the final harvest, soils were removed from the mesocosms using sterile tools, transferred to plastic bags, and homogenised by mixing. Samples were stored at 4 °C before subsampling and drying for SOC analysis. A separate subsample was stored at −20 °C until DNA extraction for characterization of microbial community.

### Soil organic carbon

For SOC determination, soil was dried at 60 °C to constant mass and ball-milled using a Retsch MM400 mill. Aliquots of 10–12 mg finely ground soil were weighed into double foil cups, treated with 50 µL of 10% HCl to remove inorganic carbon, and dried at 40 °C for 6 h. SOC concentration was then determined using a Carlo Erba NA 2500 elemental analyser (CE Instruments, UK).

Total SOC mass in each mesocosm was calculated from SOC concentration and dry soil mass. For the initial C pool, SOC contained in the 9 g γ-irradiated recipient soil was combined with SOC introduced in the 0.3 g fresh donor soil and DOC from donor soil washes used for inoculation. SOC change over the incubation was then calculated from the difference between final and initial SOC pools.

For relative changes, the log_2_ fold change was calculated as:

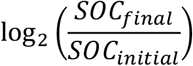

such that a value of 0 indicates no net change, negative values indicate SOC loss, and positive values indicate SOC gain.

### Amino sugar analysis and necromass estimation

Soil amino sugars were extracted and quantified following a previously published method (Salas et al., 2023) with minor modifications. Aliquots of 0.05–0.15 g air-dried soil, corresponding to approximately 4 mg soil C, were hydrolysed in 10 mL 6 M HCl at 105 °C for 8 h. Hydrolysates were filtered through 25-mm syringe filters fitted with 0.22-µm cellulose acetate membranes (VWR International, Vienna, Austria) and dried under a gentle N_2_stream. Dried residues were redissolved in 12 mL Milli-Q water, and the pH was adjusted to 6.6–6.8 with 1 M KOH. Samples were centrifuged at 2,000 × *g* for 15 min to remove precipitated metal hydroxides, primarily Fe and Al, and associated co-precipitated interferents. The supernatants were freeze-dried, redissolved in 10 mL methanol, and centrifuged again at 2,000 × *g* for 15 min to remove residual salts. The resulting supernatants were dried under a gentle N_2_stream and reconstituted in 1 mL Milli-Q water.

Amino sugars were derivatised prior to analysis using 1-phenyl-3-methyl-5-pyrazolone (PMP). Briefly, 100 µL of standard or hydrolysed sample was mixed with 100 µL 0.5 M PMP solution and 200 µL ammonia solution and incubated at 70 °C for 20 min. After cooling to room temperature, 200 µL formic acid was added to neutralise the reaction mixture. Excess PMP was removed by liquid–liquid extraction with 1 mL chloroform and 1 mL Milli-Q water. The aqueous phase containing the derivatised amino sugars was collected, filtered, and analysed using an Ultimate 3000 UPLC system (Thermo Fisher Scientific, Bremen, Germany) coupled to a Q Exactive Orbitrap high-resolution mass spectrometer equipped with a heated electrospray ionisation source.

Chromatographic separation was performed using a Waters AccQ.Tag Ultra C18 column (2.1 × 100 mm, 1.7 µm particle size; Waters, Milford, MA, USA) equipped with an ACQUITY in-line guard filter (2.1 mm, 0.2 µm). The column was maintained at 35 °C. The Q Exactive Orbitrap was operated in positive-ion full-scan mode over an *m/z* range of 150–1000.

Glucosamine (GluN) and muramic acid (MurA) were quantified as biomarkers used to estimate microbial necromass. MurA is characteristic of bacterial peptidoglycan and was therefore used as the bacterial-specific biomarker, whereas GluN occurs in both fungal and bacterial cell walls. Fungal necromass C (FNC) and bacterial necromass C (BNC) were calculated following Liang et al. (2019), accounting for the bacterial contribution to total GluN before estimating fungal-derived GluN:

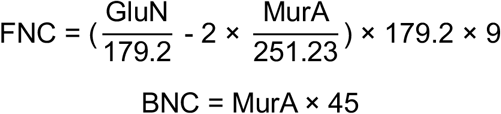

The bacterial contribution to total GluN was estimated from MurA and subtracted from total GluN to obtain fungal-derived GluN. Fungal-derived GluN and MurA were converted to fungal and bacterial necromass C using conversion factors of 9 and 45, respectively (Appuhn and Joergensen, 2006).

### DNA extractions and amplicon sequencing

DNA was extracted from 0.25 g fresh soil collected at day 229 using a DNeasy PowerSoil Pro Kit (Qiagen) according to the manufacturer’s instructions. DNA quality was assessed using a NanoDrop spectrophotometer, and DNA concentration in the final eluate was quantified using a Qubit fluorometer. DNA was also extracted from 0.25 g of dry γ-irradiated soil collected at day 0. Total extracted DNA was converted to an estimate of microbial biomass C using a conversion factor of 10.9 (Spohn et al., 2016; Malik et al., 2018).

Amplicon libraries were generated using a two-step PCR approach using Illumina Nextera-tagged primers targeting the V4–V5 region of the prokaryotic 16S rRNA gene using primers V8f ATAACAGGTCTGTGATGCCCT and v9r CCTTCYGCAGGTTCACCTAC (Bradley et al.,2016), and the fungal ITS2 region using primers ITS7f GTGARTCATCGAATCTTTG and ITS4r TCCTCCGCTTATTGATATGC (Ihrmark et al., 2012). Each primer was modified at the 5′ end with Illumina pre-adapter and Nextera sequencing-primer sequences.

Amplicons were generated using Q5 high-fidelity DNA polymerase (New England Biolabs). Following an initial denaturation at 95 °C for 2 min, PCR consisted of 30 cycles of denaturation at 95 °C for 15 s, annealing at 55 °C, 60°C, 52 °C (16S, 18S, ITS respectively) for 30 seconds, and extension at 72 °C for 30 s, followed by a final extension at 72 °C for 10 min. PCR products were purified using MultiScreenPCR filter plates (Merck) according to the manufacturer’s instructions.

MiSeq adapters and 8nt dual-indexing barcode sequences (Kozich et al., 2013) were added in a second PCR. Following initial denaturation at 95 °C for 2 min, amplification consisted of eight cycles of denaturation at 95 °C for 15 s, annealing at 55 °C for 30 s, and extension at 72 °C for 30 s, repeated for 8 cycles followed by a final 10-min extension at 72 °C.

Amplicon sizes were assessed using an Agilent 2200 TapeStation. Libraries were normalised using the NGS Normalization Kit (Norgen Biotek) and quantified using the Qubit dsDNA HS Assay Kit (Thermo Fisher Scientific). Pooled libraries were further purified by gel extraction (QIAquick, Qiagen), supplemented with 7.5% Illumina PhiX, and prepared for sequencing according to the manufacturer’s protocol. Each amplicon library was sequenced separately on an Illumina MiSeq platform using v3 600-cycle chemistry.

### Amplicon sequencing data analysis

Demultiplexed Illumina sequences were processed in R using DADA2 (Callahan et al., 2016) for quality filtering, denoising, merging, and inference of amplicon sequence variants (ASVs). Primer sequences were removed using trimLeft, and forward and reverse reads were truncated. Filtering parameters were set to a maximum number of ambiguous bases (maxN) of 5 and maximum expected errors (maxEE) of 10 for both forward and reverse reads.

Filtered reads were dereplicated, and ASVs were inferred using the DADA2 sequence-variant inference algorithm. Forward and reverse reads were merged using mergePairs, and chimeric sequences were removed using removeBimeraDenovo with default settings. Sequence tables were then constructed from the resulting ASVs.

Taxonomic assignment was performed using the assignTaxonomy function and the RDP naïve Bayesian classifier approach (Wang et al., 2007). SILVA v138.1 (Quast et al., 2013) and UNITE release 25.07.23 (Abarenkov et al., 2024) were used as reference databases for 16S rRNA gene and ITS amplicons, respectively.

### Statical analysis and visualisations

Statistical analyses were performed in R 2023.3.0 (R Core team, 2016) and figures were generated using ggplot2 (Wickham H., 2016). For microbial community analyses, sequence libraries were rarefied to a common sequencing depth after removal of samples with zero or exceptionally low read counts. Samples with zero or low counts were identified using Tukey’s outlier method (Tukey, 1977), using the vegan package in R (Oksanen et al., 2016). Rarefaction depths were 6,236 reads for 16S rRNA gene datasets and 12,335 reads for ITS datasets. β-diversity was analysed separately for each site based on Bray Curtis dissimilarity using non-metric multidimensional scaling (NMDS) and Permutational Multivariate Analysis of Variance (PERMANOVA) implemented with the adonis2 function (n = 9,999 permutations) in the vegan package. PERMANOVA models tested the effects of soil properties, inoculum source, and their interaction. ASV abundances were converted to relative abundances before ordination.

Alpha diversity was calculated from rarefied, untransformed ASV count data using the Shannon diversity index in the vegan package in R. Effects of soil properties, inoculum source, and their interaction were tested separately for each site using two-way ANOVA followed by Tukey HSD comparisons. The proportions of unique and shared ASVs among treatments were assessed using the microbiome (Lahti and Shetty, 2017) and eulerr (Larsson, 2024) packages and visualised using ggvenn package (Yan, 2025) in ggplot2.

Taxonomic, environmental, and sample-count data were integrated using the aligner() function. Samples with zero or low sequence counts were excluded as described above, and sequencing depth was standardised to 2,000 reads for indicator-taxon analysis. Dufrêne– Legendre indicator-species analysis was performed using the indval() function in the labdsv package (Roberts, 2019). Indicator ASVs associated with grassland or cropland were identified using only soils inoculated with their corresponding microbiome, with significance defined as *P* < 0.05. Their relative abundances were subsequently visualised across all treatments using the custom ngs2gel() function, which produces an electrophoresis-gel-like representation of ASV occurrence and abundance. ASVs were ordered taxonomically and band thickness scaled to relative abundance. Indicator taxa were highlighted according to grassland or cropland association, with phylum- and order-level taxonomy annotated.

We tested for statistical difference in necromass and SOC across treatment combinations for each site separately. Linear mixed-effects were used to test the effects of soil properties (grassland vs. cropland), inoculum source (grassland vs. cropland), and their interaction on microbial biomass C, fungal and bacterial necromass, final SOC, and SOC change. Models were fitted with the lm function in R with the model structure: Soil*Inoculum. Effects of soil properties, inoculum source, and their interaction were also tested using two-way ANOVA followed by Tukey HSD comparisons for post hoc pairwise analysis. Model residuals were inspected to assess normality and homoscedasticity, and response variables were transformed where necessary to meet model assumptions.

## CRediT authorship contribution statement

Conceptualization: AAM

Formal analysis: LC, TL, TG, AAM

Methodology: LC, TL, TG

Investigation: LC, TL, TG, AAM

Project administration: AAM, CG-R

Resources: WW, EP, CG-R, AAM

Validation: LC

Data curation: LC

Visualization: LC, TG, RG, AAM

Funding acquisition: LC, AAM, CG-R, WW

Supervision: AAM, CG-R

Writing – original draft: LC, AAM

Writing – review & editing: All authors

## Acknowledgements

This work was funded by a NERC-sponsored Daphne Jackson Trust Fellowship awarded to LC. AAM was supported by University of Edinburgh Chancellor’s Fellowship in Climate and Environmental Sustainability. C.G.-R was supported by a Royal Society University Research Fellowship (URF150571). We also wish to thank Georgina Bray at RSPB Knapwell, Cambridgeshire and Paul Hill at The Earth Trust, Oxfordshire for permission for soil sampling. For technical support, we thank Gafaru Sumaila, Alfonso Hernandez Molina, Nucia Jamshidi, David Coutts, Michael McGibbon and David Galloway at the University of Aberdeen, and Gillian Martin at The James Hutton Institute.

