## Supplementary figures and tables for "Soil microenvironment and microbial community composition jointly regulate carbon accrual in agricultural soils"

**Table S1:** Physico-chemical properties of soils from the site-land use treatments given as mean values (± standard error).

| Soil properties | Site 1 grassland | Site 1 cropland | Site 2 grassland | Site 2 cropland |
|---|---|---|---|---|
| Data from this study, sampled in 2022 | | | | |
| Soil moisture (%) | 37.49 (± 1.25) | 16.98 (± 0.52) | 30.06 (± 1.19) | 22.85 (± 3.23) |
| Soil pH (in water) | 7.70 (± 0.05) | 7.72 (± 0.03) | 7.40 (± 0.04) | 7.59 (± 0.14) |
| Soil organic carbon (%) | 7.56 (± 0.62) | 2.75 (± 0.05) | 6.88 (± 0.01) | 2.54 (± 0.06) |
| Data from earlier study, sampled in 2017 (Malik et al., 2016) | | | | |
| Soil moisture (%) | 41.76 (± 3.14) | 27.42 (± 3.46) | 35.18 (± 2.71) | 23.60 (± 0.31) |
| Soil pH (in water) | 7.60 (± 0.14) | 7.94 (± 0.02) | 7.47 (± 0.08) | 7.67 (± 0.03) |
| Soil organic carbon (%) | 7.38 (± 1.04) | 3.98 (± 0.1) | 6.29 (± 1.1) | 2.06 (± 0.05) |
| Soil organic nitrogen (%) | 0.72 (± 0.11) | 0.34 (± 0.02) | 0.57 (± 0.09) | 0.20 (± 0.01) |
| SOM C:N ratio | 10.22 (± 0.34) | 11.84 (± 0.25) | 11.08 (± 0.09) | 10.15 (± 0.01) |
| Clay % | 33 | 41 | 31 | 31 |
| Clay index from FTIR | 0.22 (± 0.05) | 0.27 (± 0.02) | 0.13 (± 0.03) | 0.10 (± 0.002) |

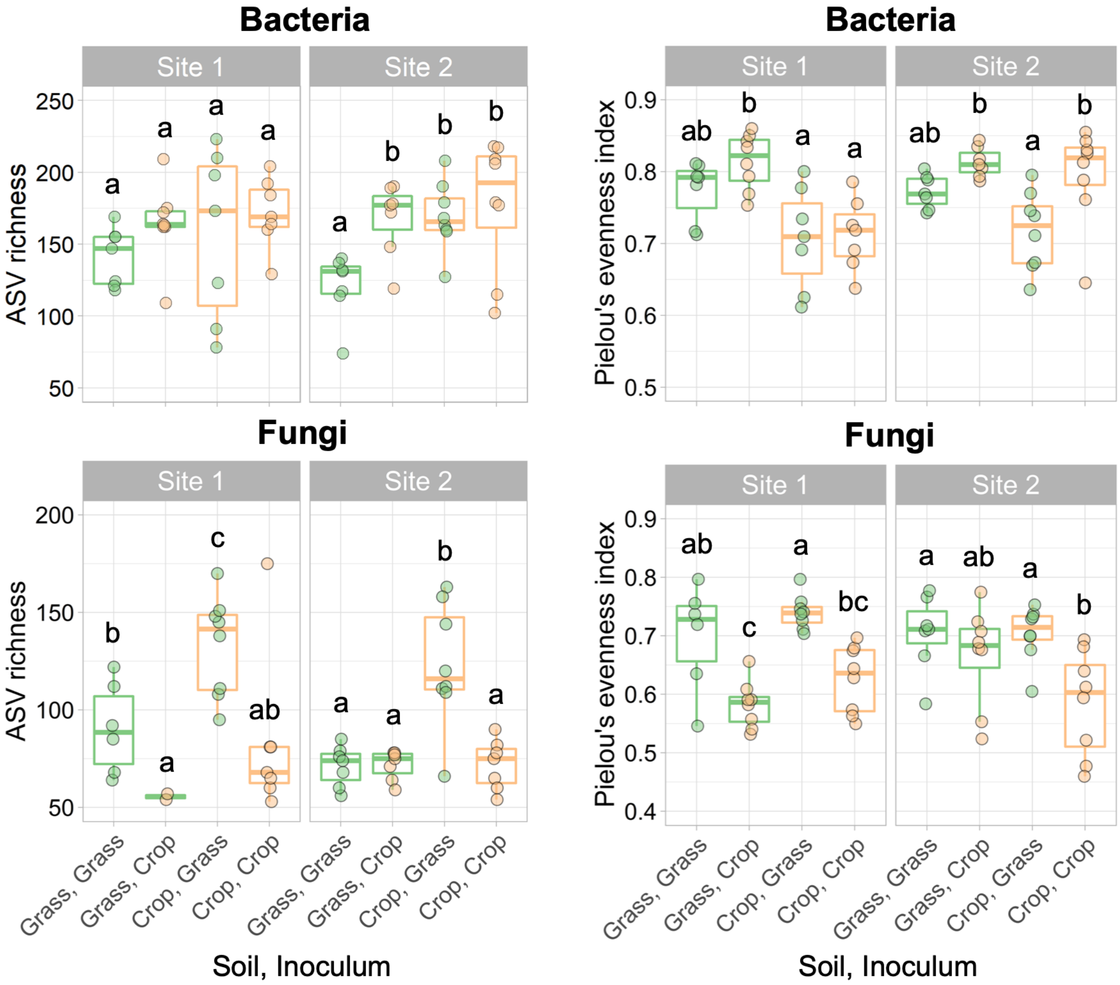


**Fig. S1. Alpha diversity of assembled prokaryotic and fungal communities.** ASV richness of prokaryotes (a) and fungi (c), and Pielou's evenness of prokaryotes (b) and fungi (d), at day 229 for Site 1 and Site 2 mesocosms. Boxplots show cropland and grassland recipient soils, with individual points indicating cropland- or grassland-derived inocula. Letters indicate pairwise Tukey HSD differences at $P < 0.05$, analysed separately for each site; treatments sharing a letter did not differ significantly.

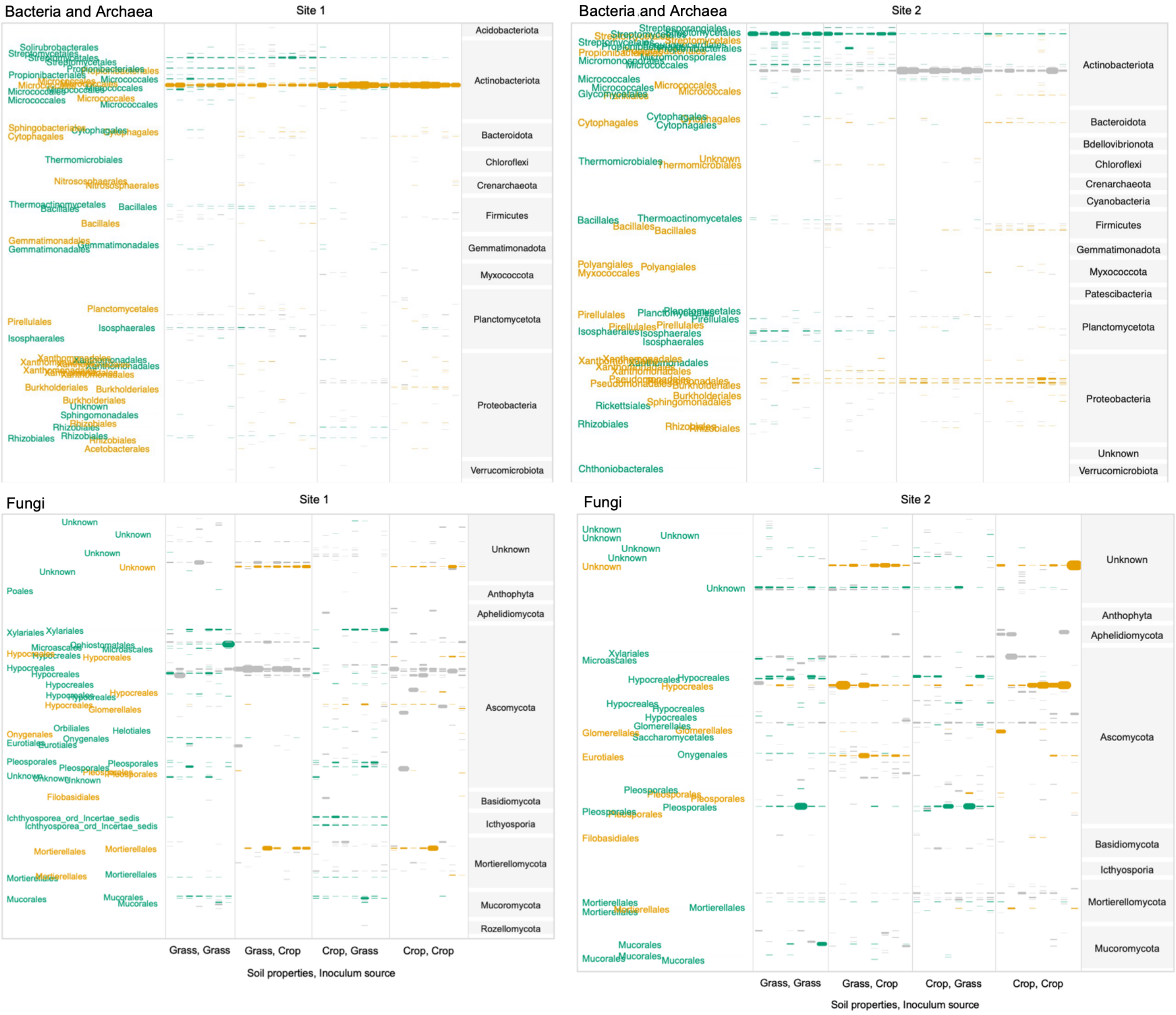


**Fig. S2. Distribution of indicator taxa across treatments.** Prokaryotic and fungal indicator ASVs were identified using Dufrêne–Legendre indicator-species analysis. Grassland and cropland recipient soils inoculated with their corresponding microbial communities were used to identify significant indicator ASVs. Individual bands represent ASVs, with grassland-associated taxa highlighted in green and cropland-associated taxa in orange. ASVs are ordered taxonomically, and band thickness represents relative abundance across treatments.

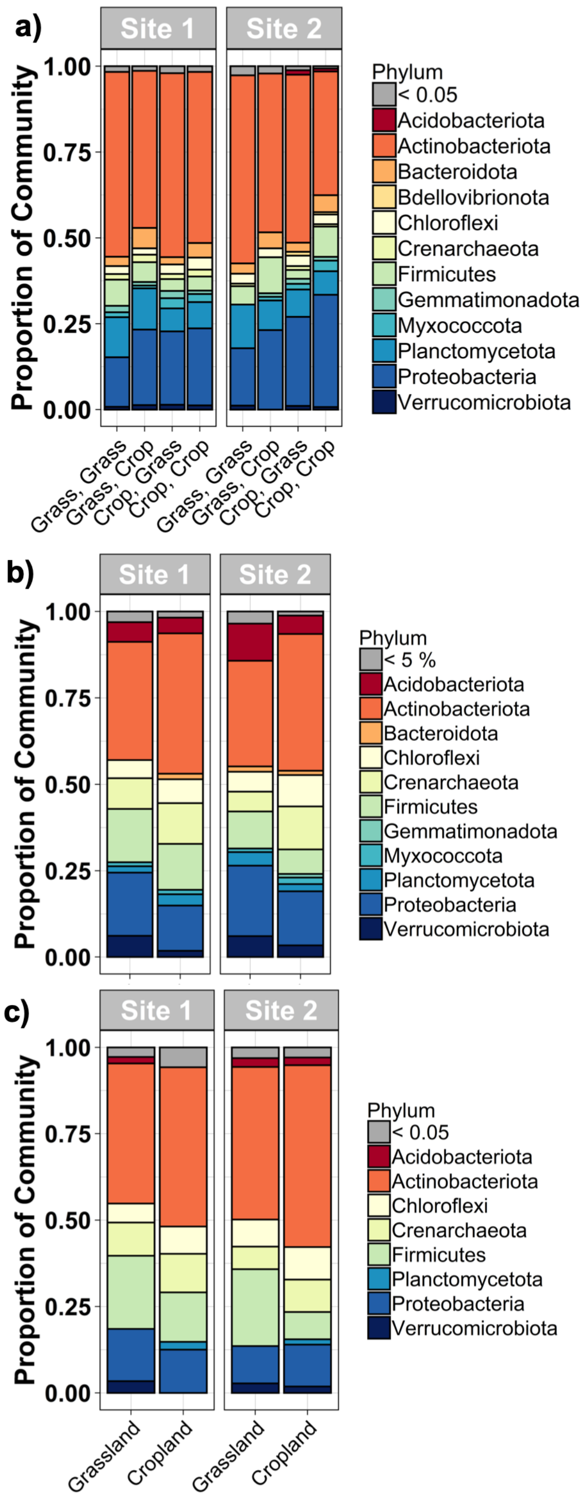


**Fig. S3. Relative abundance of prokaryotic phyla across experimental and reference soils.** Relative abundances of prokaryotic phyla at (a) day 229 in the experimental mesocosms, (b) non-sterilised soils, and (c) sterilised soils from Site 1 and Site 2. Treatment labels in (a) denote recipient soil followed by inoculum source: Grass, Grass: grassland soil with grassland inoculum; Grass, Crop: grassland soil with cropland inoculum; Crop, Grass: cropland soil with grassland inoculum; and Crop, Crop: cropland soil with cropland inoculum. Panels (b) and (c) show the corresponding grassland and cropland soils.

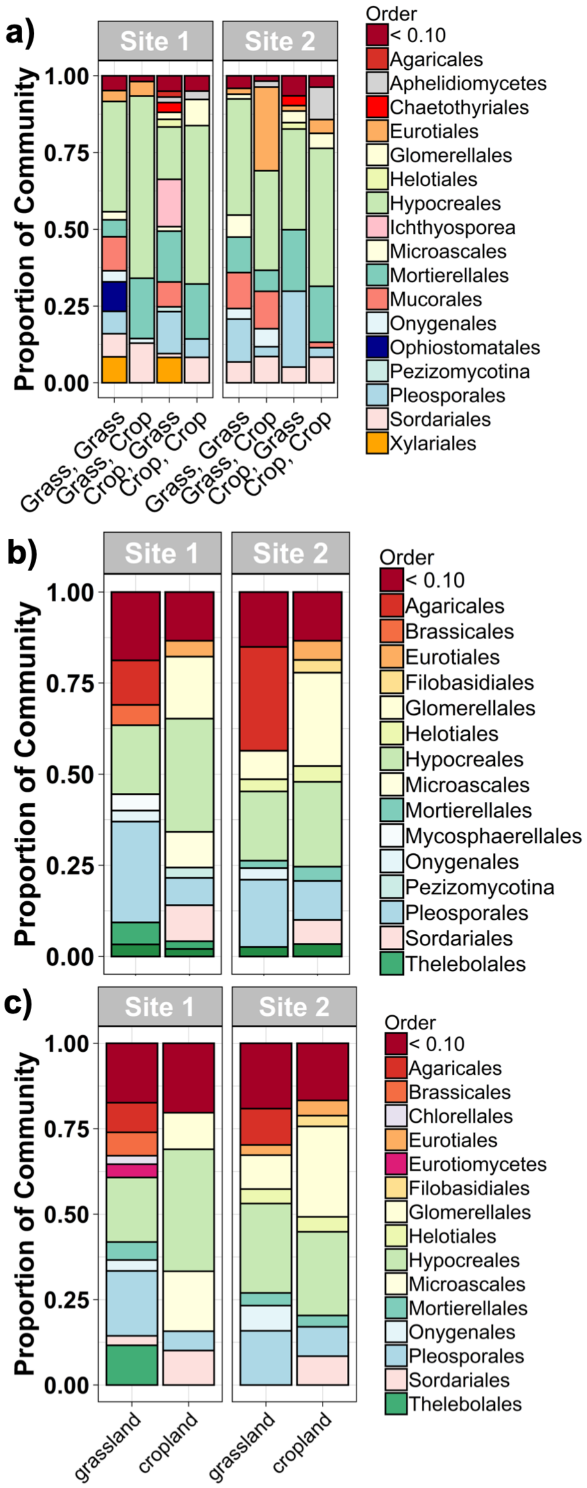


**Fig. S4. Relative abundance of fungal phyla across experimental and reference soils.** Relative abundances of fungal phyla at (a) day 229 in the experimental mesocosms, (b) non-sterilised soils, and (c) sterilised soils from Site 1 and Site 2. Treatment labels in (a) denote recipient soil followed by inoculum source: Grass, Grass: grassland soil with grassland inoculum; Grass, Crop: grassland soil with cropland inoculum; Crop, Grass: cropland soil with grassland inoculum; and Crop, Crop: cropland soil with cropland inoculum. Panels (b) and (c) show the corresponding grassland and cropland soils.

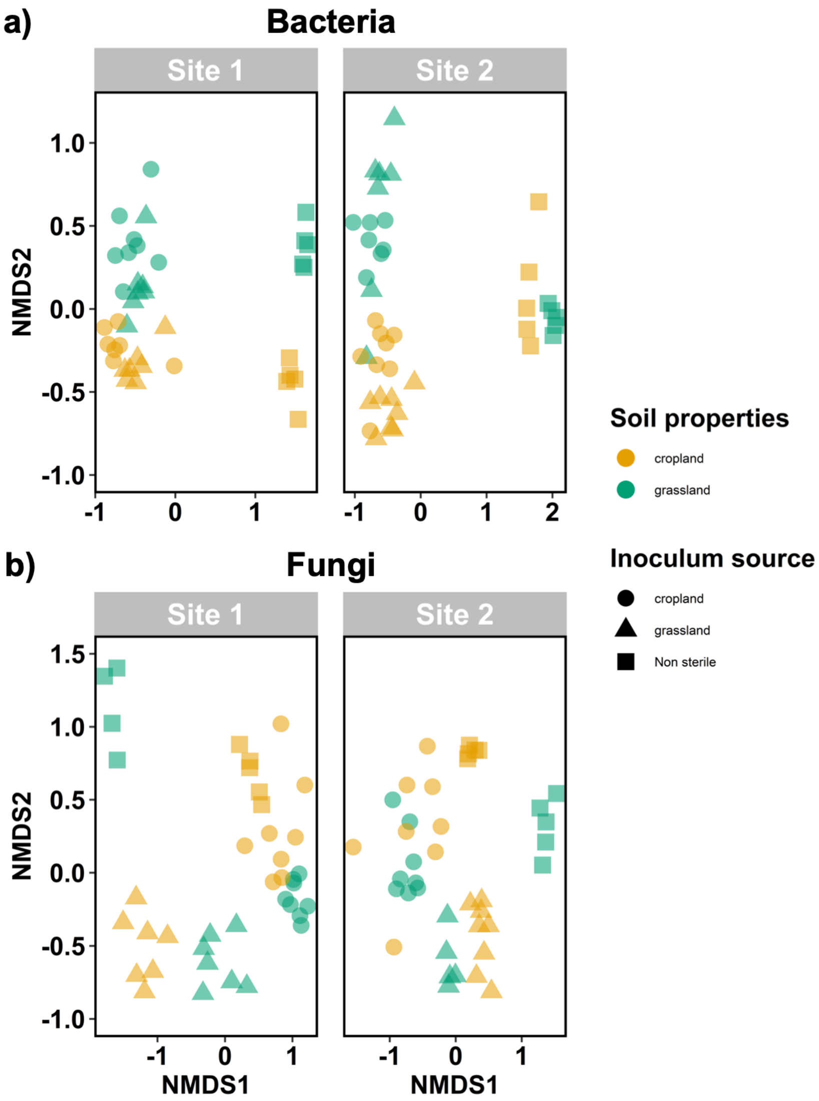


**Fig. S5. Comparison of assembled mesocosm communities with non-sterilised soil communities.** NMDS ordinations of (a) prokaryotic ASVs derived from 16S rRNA gene sequencing and (b) fungal ASVs derived from ITS sequencing, including experimental mesocosms from both sites and non-sterilised soils. Point colour denotes recipient soil (grassland, green; cropland, orange), whereas point shape denotes inoculum source (grassland, triangles; cropland, circles); non-sterilised field soils are shown as squares.

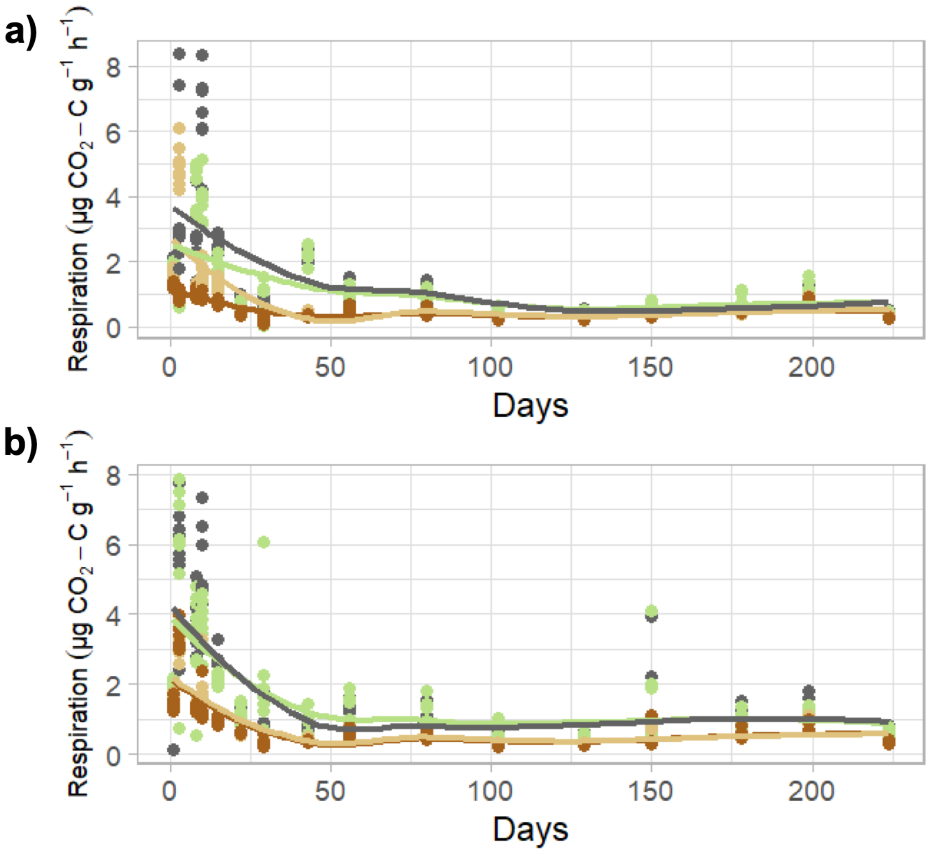


**Fig. S6. Soil respiration over the eight-month incubation.** $CO_2$-C release from mesocosms containing (a) Site 1 and (b) Site 2 soils. Lines show smoothed treatment means (*n* = 8): cropland soil with cropland inoculum (dark brown), cropland soil with grassland inoculum (beige), grassland soil with grassland inoculum (dark green), and grassland soil with cropland inoculum (light green).

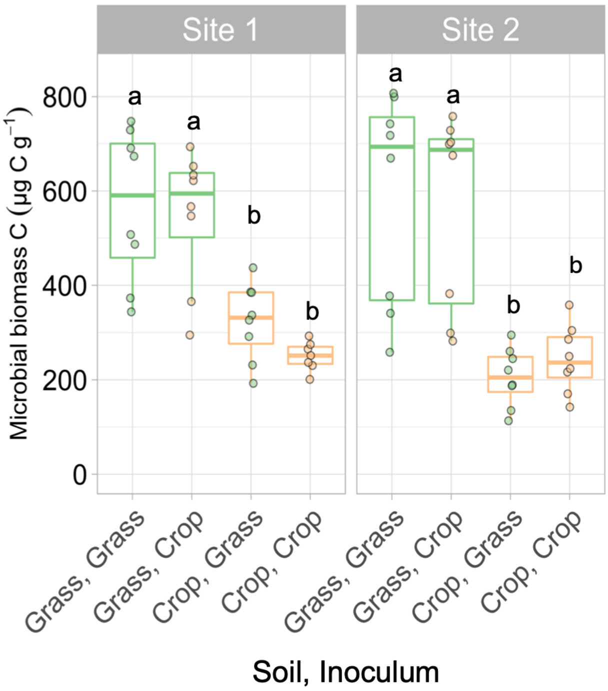


**Fig. S7. Microbial biomass C estimated from DNA yield at the end of the experiment.** Microbial biomass C was estimated from DNA yield at day 229. Letters indicate pairwise Tukey HSD differences at $P < 0.05$, analysed separately for each site; treatments sharing a letter did not differ significantly. Thick horizontal lines indicate medians, boxes represent interquartile ranges, and whiskers extend to 1.5 times the interquartile range.
